# New Record of Four Thrips Species Infesting Flower Buds of *Magnolia champaca* in India

**DOI:** 10.1101/2024.10.30.621126

**Authors:** R.R. Rachana, D.M. Firake, Kaomud Tyagi, K.V. Prasad, B. Amarendra

## Abstract

The golden-flowered, highly fragrant local variety of *Magnolia champaca*, known as ‘Sonchafa,’ is very popular loose flower crop in Maharashtra and adjoining states of India. In response to the rising thrips problems in Sonchafa (*M. champaca*) fields leading to significant economic losses, we looked into the thrips’ species composition infesting the plant in the Western Maharashtra region of India. Four thrips species were newly recorded as pests, including *Psephenothrips machili* (Moulton), *Thrips florum* Schmutz, *Thrips hawaiiensis* (Morgan) and *Thrips orientalis* (Bagnall). Among these, *P. machili* was identified as the dominant species, responsible for 63% to 71% of bud damage across various fields. This study also marks the first report of an anthophilous host association of *P. machili* with *M. champaca*, causing significant economic damage. The observed damage symptoms include, the typical distortion of flower tissues and buds, black spots on buds were ascribed to *P. machili*. The other three species were found in groups and caused silvery streaks on early-stage buds. The diagnostic characteristics of all the reported species are provided here, accompanied by photomicrographs.

## Introduction

The champaka tree, *Magnolia champaca* (L.) Baill. ex Pierre (=*Michelia champaca* L.) (Magnoliaceae), is an evergreen species valued for its commercial, industrial, and aesthetic uses. The golden-flowered, highly fragrant local variety of *M. champaca*, known as ‘Sonchafa,’ is popular loose flower crop in Maharashtra and adjoining states of India. Its exceptional fragrance and distinctive golden blossoms have made it a favoured choice among floriculture farmers in this region. The Sonchafa shrub blooms year-round, producing 150 to 200 flowers daily per shrub [1] and each plant can generate income of Rs. 250-300 per day, with price rising to Rs. 1400 during festive seasons [4]. Hence, Sonchafa has become an important source of livelihood and prosperity for many farmers in Maharashtra and adjoining states.

Globally, several pests and diseases are associated with *M. champaca*, and some of them cause significant damage to this plant [6]. In India, the insect pests recorded feeding on *M. champaka* include *Liothrips* (*Liothrips*) *champakae* (Ramakrishna & Margabandhu) (synonym: *Rynchothrips champakae*) [8]; *Urotylis punctigera* [12]; *Podothrips* sp. [9]; *U. parapunctigera* [7] and *Graphium doson* [5].

Accurate pest species identification is the foremost step towards developing robust management practices. Vigilant observation of various species of pest on an economically important crop allows investigators to evaluate precise shifts in pest species complexes, mainly for sucking insects such as thrips which can be moved afar naturally or through accidental human aid. To date, only two thrips species, *L. champakae* and *Podothrips* sp., have been reported infesting *M. champaca* in India, both species causing damage to the leaves, twigs, and stems of the plant.

The objectives of the current study is to examine and catalogue the thrips’ species composition damaging flower buds of *M. champaca* in Maharashtra, India, to depict the symptoms of damage, and to illustrate the significant diagnostic characters of different species infesting various plant parts of the *M. champaca*. According to the primary scientific literature, this study is the initial footstep to understand the thrips species infesting flowers, the economic yield of *M. champaca*.

## Materials and methods

Surveys were conducted in Sonchafa, *M. champaca* plantations during the peak flowering period (August-September) at Purandhar (Pune), Kudal (Sindhudurg), Shahapur (Thane), and Vasai-Virar (Palghar) in Western Maharashtra. Five fields in each area were examined for various insect pests causing significant damage to *M. champaca* plantations. During the surveys, severe incidence of thrips on the flower buds of *M. champaca* were observed at several locations. The per cent damage caused by thrips was determined by comparing the number of thrips-infested flower buds to the total number of flower buds examined in each field.

Collected thrips were kept in vials with AGA medium for supplementary studies. The specimens were mounted on microslides using Canada balsam, examined through an Olympus BX 51 microscope, imaged with Olympus DP 23 camera mounted on the microscope. The species level identities of the specimens were determined using the diagnostic keys [3, 14]. Voucher specimens were deposited in the National Insect Museum (NIM) at the Indian Council of Agricultural Research – National Bureau of Agricultural Insect Resources (ICAR–NBAIR), Bengaluru, India and Zoological Survey of India (ZSI), Kolkata, India.

## Results

### Thrips infestation to *M. champaka* flower buds in farmer’s fields in Western Maharashtra

Four thrips species were found infesting *M. champaka* flower buds in farmer fields in Western Maharashtra region of India *viz*., *Thrips florum* Schmutz, *Thrips hawaiiensis* (Morgan), *Thrips orientalis* (Bagnall) (Thysanoptera: Thripidae) and *Psephenothrips machili* (Moulton) (Thysanoptera: Phlaeothripidae). Even though all four species were found together as mixed populations in the field, *P. machili* was the most abundant. Both larvae and adults of *P. machili* were observed feeding on *M. champaca* buds, causing distortion of flower tissues and buds, which led to unattractive appearance of the buds and flowers. Severe infestation resulted in black spots on buds and fully deformed buds, rendering them unmarketable (Fig. 1). Although less abundant, the other three thrips species were found in groups and caused silver streaks on early-stage buds and brown necrotic spots on fully developed buds (Fig. 2).

**Fig. 1.**
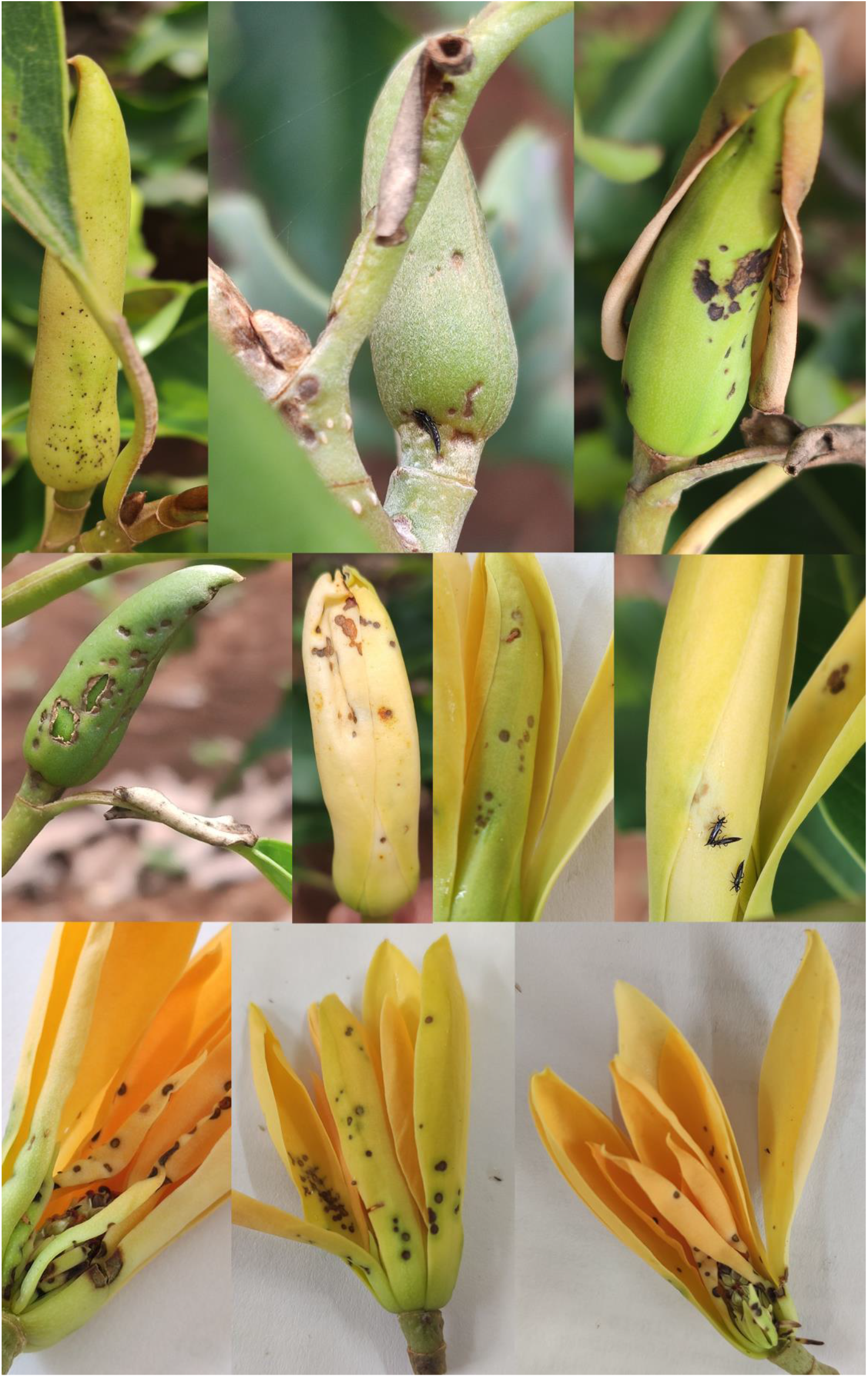
Damage symptoms caused by *Psephenothrips machili*

**Fig. 2.**
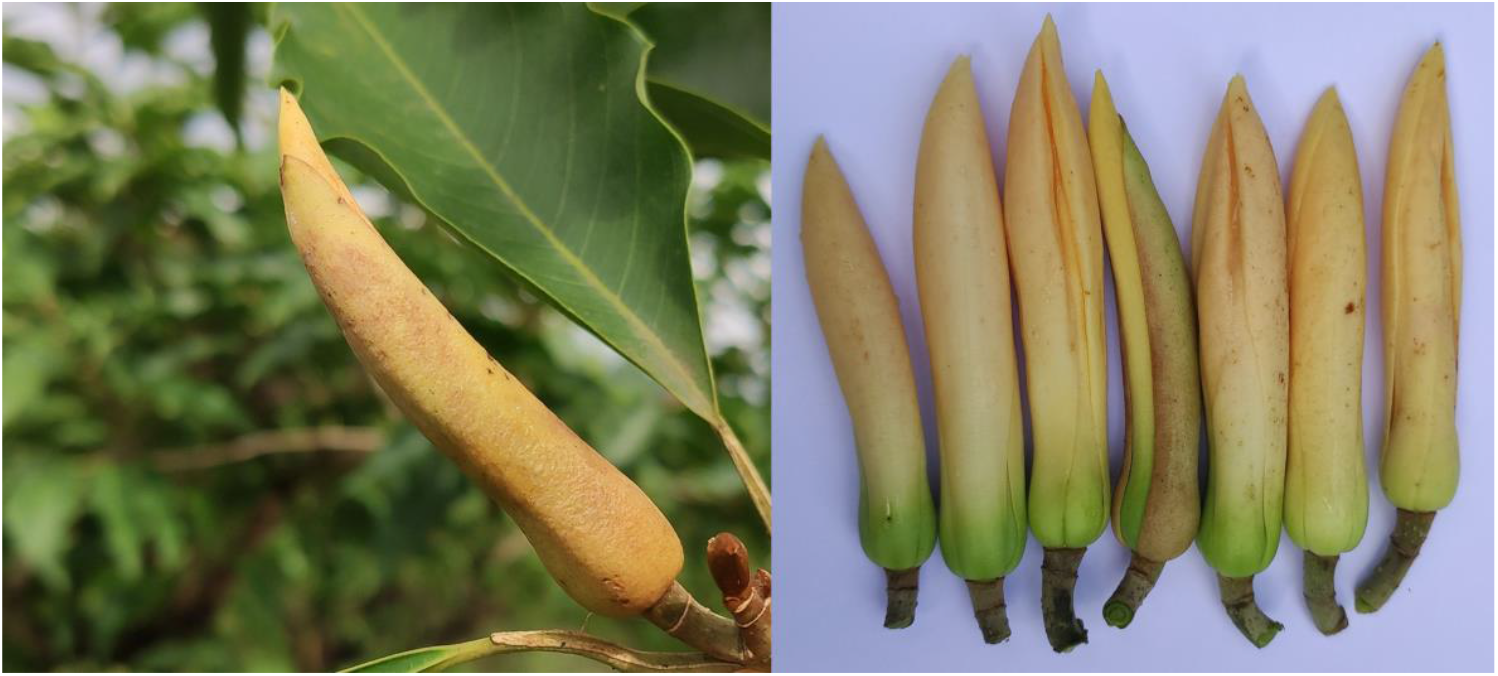
Damage symptoms caused by *Thrips florum, Thrips hawaiiensis* and *Thrips orientalis*

The highest infestation of *P. machili* was found in Sonchafa (*M. champaca*) fields at Kodit village (Purandar, Pune), Maharashtra, with infestation levels ranged from 63 to 71% across different fields. Although the incidence was not severe, a mixed infestation of *T. florum* and *T. hawaiiensis* was recorded in the sonchafa farms at Kodit village (Purandar, Pune) (3.33-6.67%); in Sahyadri farm at Sajivali (Shahaur, Thane) (2.67-3.33%); sonchafa farm at Vetal-bambarde village (Kudal, Sindhudurg) (2.33-4.67%), and in several sonchafa farms (4.33-7.67%) at Vasai-Virar area of Palghar district of Maharashtra. Infestation by *T. orientalis* was recorded only in the sonchafa farms at Kodit village (Purandar, Pune), with infestation ranged from 7.33 to 11.67 per cent.

### Diagnosis of thrips species infesting *M. champaka* flower buds

Important diagnostic characters of different thrips species infesting *M. champaca* have been presented below;

### *Psephenothrips machili* (Moulton) (Fig. 3)

Body brown including femora, tibiae and tarsi, distal half of fore tibiae pale, yellowish brown fore tarsi; dark brown antennal segments I, II, VII and VIII, distal half of segment II and extreme base of segment VII yellowish, III yellow, IV largely yellow with slightly shaded apex, V yellow with apex brown, VI brown with distal third yellow. Head longer than broad; cheeks weakly rounded, gradually narrowed towards base, a little narrowed at base. Mouth-cone elongated reaches upto the anterior margin of ferna; maxillary stylets very elongated, touching eyes, nearly touching together; maxillary bridge absent. Pronotum almost smooth, with a short median longitudinal line; five pairs of major setae blunt or very weakly expanded at apex. Metathoracic sternopleural sutures absent. Forewing with 20-26 duplicate cilia; three sub-basal wing setae weakly expanded at apex. Pelta polygonally reticulate. Abdominal tergum IX S1setae longer than S2, almost pointed or with narrowly blunt apex. Tube shorter than head, slightly constricted at apex. Anal setae shorter than tube. Male similar to female, without fore tarsal tooth. Forewing with 16-22 duplicated cilia. Glandular area on sternum VIII absent. Tube shorter than head.

**Fig. 3.**
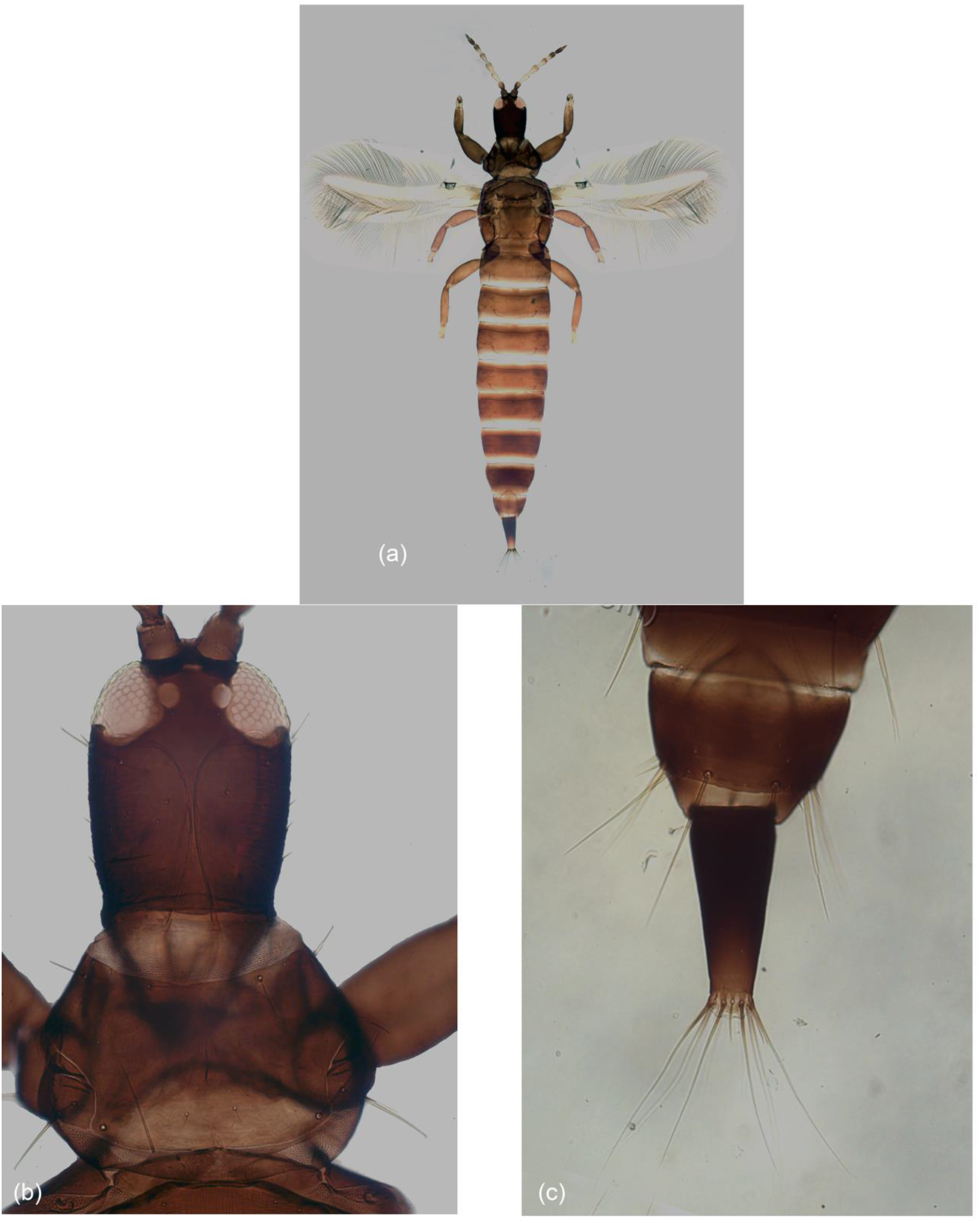
*Pseophenothrips machili* (a) Female; (b) Head and Pronotum; (c) tergites IX-X

### *Thrips florum* Schmutz (Fig. 4)

Body brown, yellow antennal segment III; brown fore wing having paler base. Antennae 7 segmented; ocellar setae III placed exterior to ocellar triangle; postocular seta I and III longer than II. Metanotum transversely striate anteriorly, widely spaced longitudinal striations posteriorly, median setae positioned nearly at anterior margin, with campaniform sensilla. First vein of forewing with 3 distal setae, clavus with terminal seta shorter than subterminal seta. Posteromarginal complete on abdominal tergite VIII. Discal setae present on abdominal sternites III–VII.

**Fig. 4.**
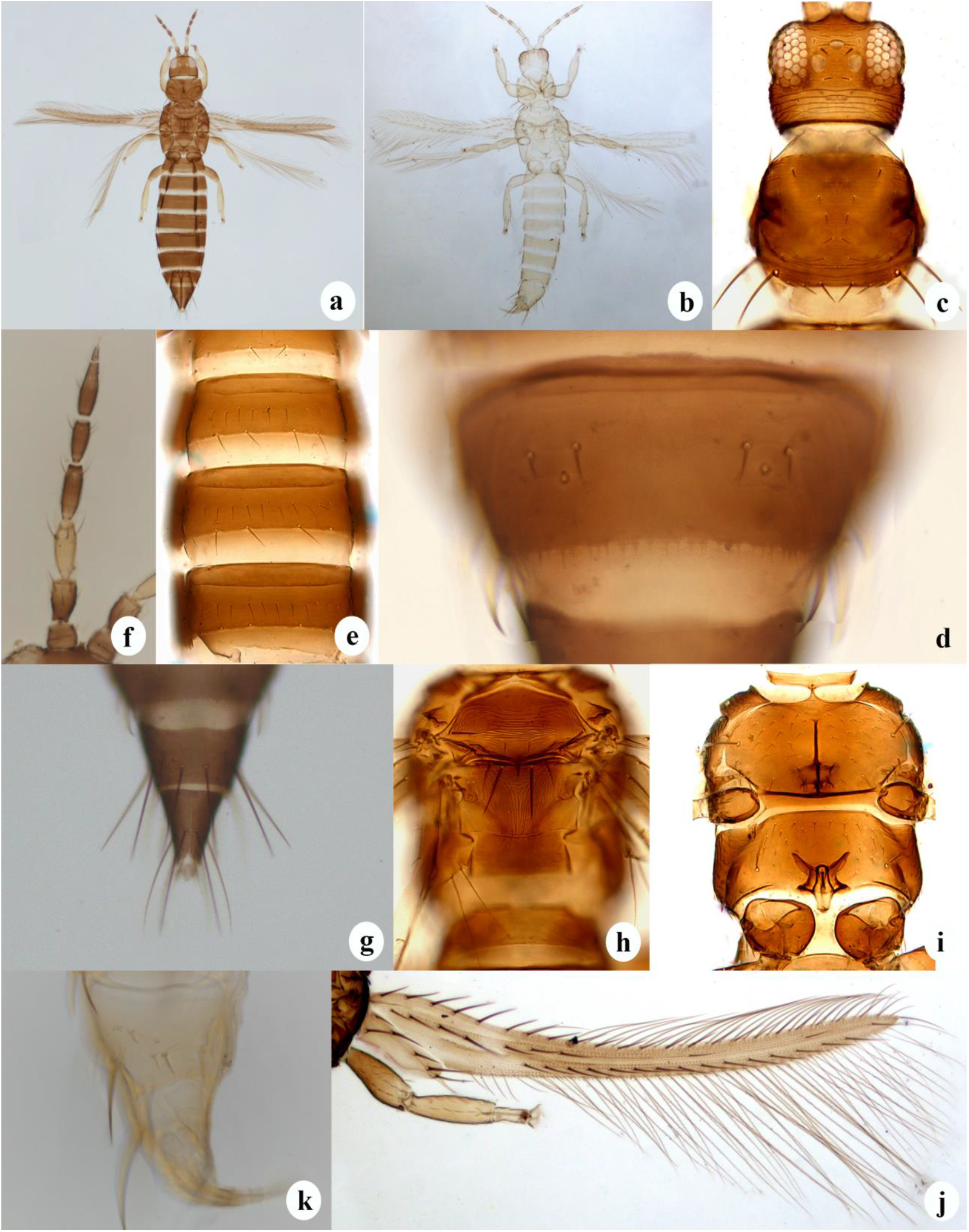
*Thrips florum* (a) Female; (b) Male; (c) Head and prothorax; (d) Abdominal tergite VIII; (e) Abdominal sternites III-VI; (f) Antenna; (g) Abdominal tergites IX-X (female); (h) Meso and metathorax; (i) Pterosterna; (j) Fore wing; (k) Abdominal tergites IX-X (male)

### *Thrips hawaiiensis* (Morgan) (Fig. 5)

Bicoloured body with orange head and thorax and brown abdomen; yellow antennal segment III; brown fore wing having paler base. Antennae 7 segmented; ocellar setae III positioned exterior to ocellar triangle; subequal postocular setae I and II. Metanotum transversely striate anteriorly, with longitudinal striations posteriorly, median setae placed at anterior margin, with campaniform sensilla. First vein of forewing with 3 distal setae, clavus with subterminal seta shorter than terminal seta. Complete posteromarginal comb on abdominal tergite VIII; discal setae present on abdominal sternites III–VII.

**Fig. 5.**
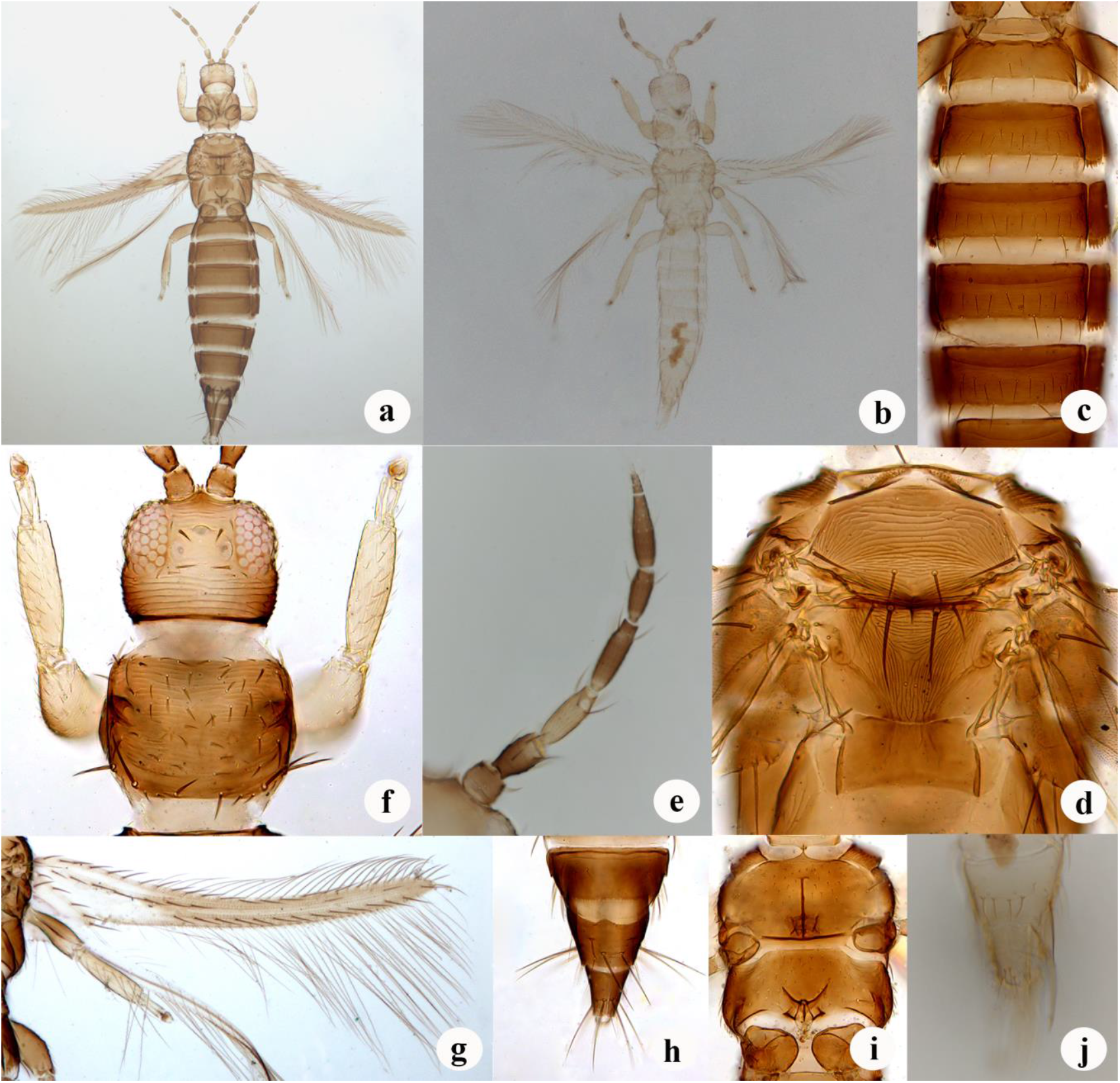
*Thrips hawaiiensis* (a) Female; (b) Male; (c) Abdominal sternites II-VI; (d) Meso and metathorax; (e) Antenna; (f) Head and prothorax; (g) Fore wing; (h) Abdominal tergites IX-X (female); (i) Pterosterna; (j) Abdominal tergites IX-X (male)

### *Thrips orientalis* (Bagnall) (Fig. 6)

Body, legs and forewings brown; antennal segment III yellow. Antennae 7-segmented; ocellar setae III positioned at lateral margins of ocellar triangle, minute postocular setae II. Metanotum reticulate with internal markings, median setae posterior to anterior margin, without campaniform sensilla. First vein of forewing with almost uniformly arranged setae. Abdominal tergite VIII without median posteromarginal comb; sternites III–VI sometimes without discal setae but generally with 1–6 setae laterally.

**Fig. 6.**
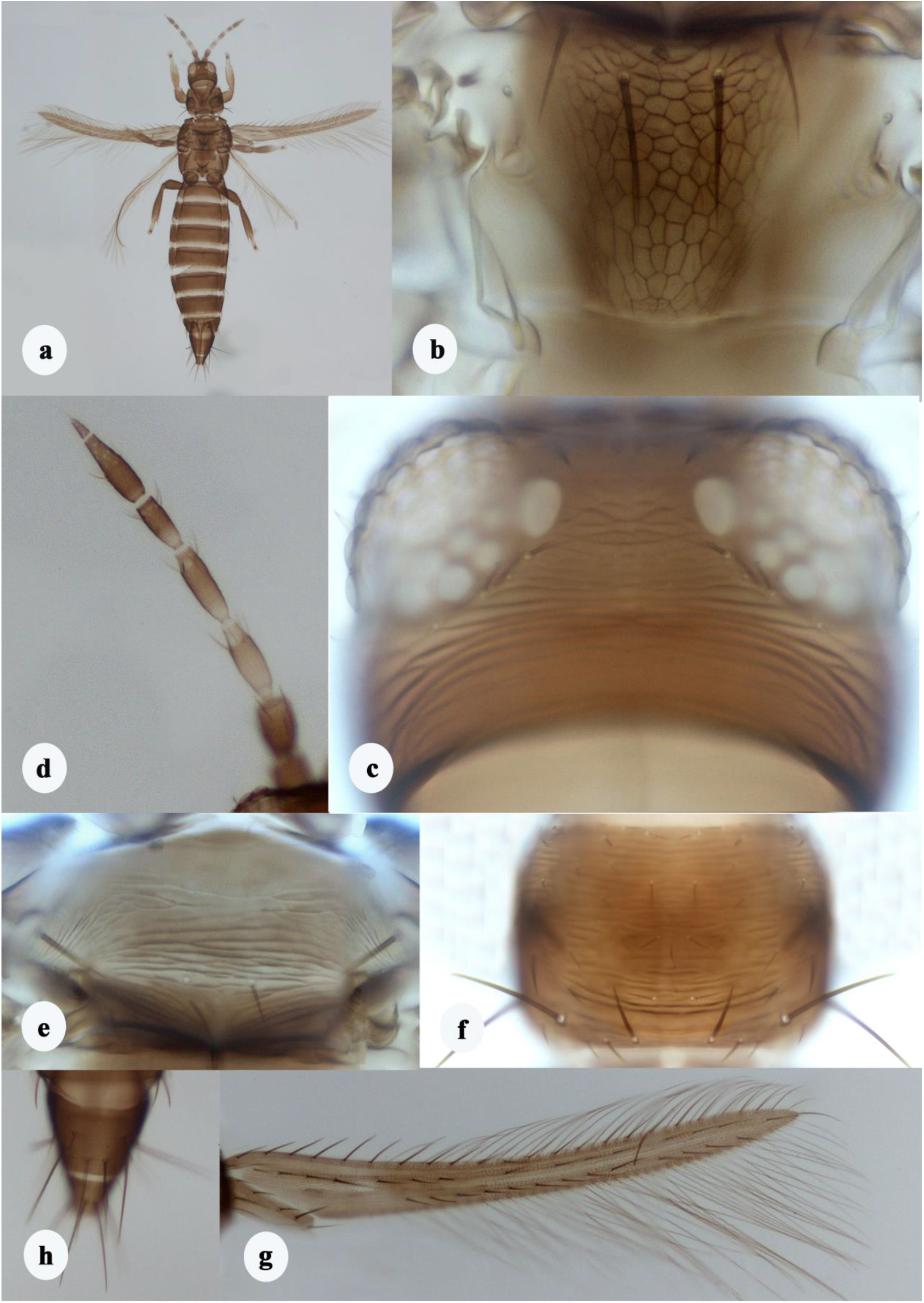
*Thrips orientalis* (a) Female; (b) Metathorax; (c) Head; (d) Antenna; (e) Mesothorax; (f) Prothorax; (g) Fore wing; (h) Female abdominal tergites IX-X

## Discussion

Globally, several insect pests are associated with *Magnolia champaca*, affecting different plant parts at various growth stages. Two species of urostylid bugs, *Urotylis punctigera* [12] and *Urotylis parapunctigera* [7], are known to cause extensive damage to *M. champaca*, sometimes leading to the death of the trees. Earlier records indicated that the thrips species, *Liothrips* (*Liothrips*) *champakae* infests the leaves of *M. champaca*, resulting in browning and death of the leaves and twigs [8]. *Podothrips* sp. has also been reported to cause stem galls in *M. champaca* [9]. Additionally, two other thrips species, *Scirtothrips citri* (Moulton) in Ireland [10] and *Tryphactothrips rutherfordi* (Bagnall) in Sri Lanka [13], have been recorded as damaging *M. champaca*.

We recorded four additional thrips species for the first time infesting *M. champaca* flower buds *viz*., *Psephenothrips machili, Thrips florum, Thrips hawaiiensis* and *Thrips orientalis*. Among them, *P. machili* was dominant species found causing extensive damage to *M. champaka* flower buds. *P. machili* was earlier found to be associated with leaves of *Machilus thunbergii* (Lauraceae) and *Ilex integra* (Aquiforiaceae) in Japan [11] and leaf galls of *Machilus macranthes* in India [2]. This study reports for the first time an anthophilous host association of *P. machili* on *M. champaca* causing severe economic damage to the flower buds of the crop and thus reducing significant market value. Two other species viz., *Thrips florum* Schmutz and *Thrips hawaiiensis* (Morgan) are polyphagous species commonly found associated with various ornamental flowers. The fourth recorded species is *Thrips orientalis* (Bagnall), which is known to infest flowering plants particularly of the genera *Jasminum, Gardenia*, and *Plumeria*. Our study exhorts growers of *M. campaca* to consider infestation of thrips on flowers as a complex of different species rather than a sole species. The photomicrographs presented here will be helpful for the easy understanding of field level damage symptoms as well as authentic identification of these thrips species.

## Acknowledgements

The funding for the institute project “Eco-friendly Pest Management in Commercial Loose Flower Crops, Project code: 45/S/IPP/12” by the Indian Council of Agricultural Research, New Delhi is duly acknowledged.

## Statements and Declarations

Authors declare that this work has not been published previously or it is not under consideration for publication elsewhere. The article’s publication is approved by all the authors and tacitly or explicitly by the responsible authorities where the work was carried out

